# HCC-1 accelerates atherosclerosis by inducing endothelial cells and macrophages pyroptosis and serves as an early diagnostic biomarker

**DOI:** 10.1101/2023.03.22.533883

**Authors:** Fan Bu, Junhui Wang, Qi Zhang, Xiaomin Lin, Ruyi Zhang, Huanlan Bai, Juanjiang Chen, Yuneng Hua, Haifang Wang, Mei Huang, Yiyi Huang, Xiumei Hu, Lei Zheng, Qian Wang

**Affiliations:** Center for Clinical Laboratory, Zhujiang Hospital, Southern Medical University, Guangzhou, 510260, People’s Republic of China; Department of Laboratory Medicine, Nanfang Hospital, Southern Medical University, Guangzhou, 510515, People’s Republic of China; Department of Hematology, Nanfang Hospital, Southern Medical University, Guangzhou, 510515, People’s Republic of China #: co-first author

**Keywords:** atherosclerosis, chemokines, inflammation, pyroptosis, endothelial cells, macrophages

## Abstract

**BACKGROUND:** HCC-1 (Hemofiltrate CC chemokine-1), a CC-type chemokine, exerts function to change intracellular calcium concentration, induce leukocyte and manipulate enzyme release especially in monocytes. It has been reported that HCC-1 could predict the persistent acute kidney injury (AKI) or suppress hepatocellular carcinoma (HCC) by modulating cell cycle and promoting apoptosis, but the effect of HCC-1 on atherosclerosis is poorly understood. Here, we aimed to clarify the function and mechanism of HCC-1 in atherosclerosis and whether it could serve as a novel biomarker for the diagnosis of atherosclerosis.

**METHODS:** HCC-1 expression in serum, atherosclerotic plaques and normal arterial tissue from patients with atherosclerosis and control group was assessed by enzyme-linked immunosorbent assay, immunohistochemistry and con-focal microscope. The atherosclerotic model of HCC-1 overexpressing mice and control mice were generated by infection of AAV9-HCC-1 on an ApoE^−/−^ background. Cell adhesion, polarization and pyroptosis were evaluated in vitro. The relationship between HCC-1 concentration in serum and atherosclerosis was analyzed in patients with atherosclerosis.

**RESULTS:** HCC-1 expression increased in patients with atherosclerosis both in serum and atherosclerotic plaque. HCC-1 overexpression mice had an enhancement in macrophage accumulation in plaque, higher levels of inflammatory factors, increased pyroptotic rate in ECs and Macrophages in plaque and decreased atherosclerotic plaque stability. In vitro, HCC-1 promoted monocytes to adhere to endothelial cells and M1 polarization, induced inflammation and pyroptosis both in ECs and Macrophages.

**CONCLUSIONS:** HCC-1 expression markedly increased in patients with atherosclerosis and HCC-1 overexpression accelerated atherosclerotic burden via an enhancement in monocytes recruitment, M1 polarization and pyroptosis both in ECs and Macrophages. Our findings suggested that HCC-1 may serve as an early biomarker for the diagnosis of atherosclerosis, with the capacity to reflect the degree of stenosis.

**GRAPHIC ABSTRACT:** A graphic abstract is available for this article.

## Nonstandard Abbreviations and Acronyms

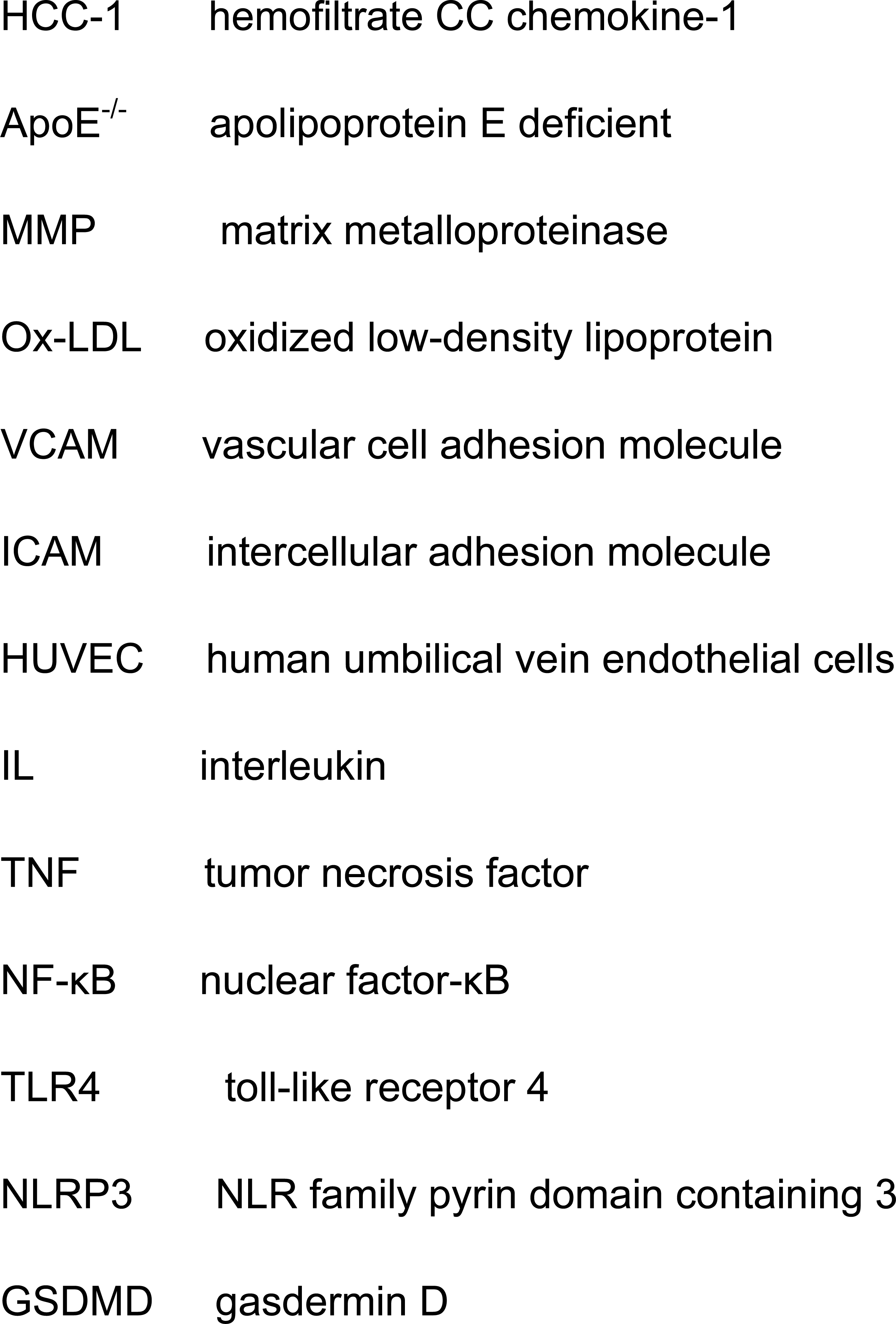

## INTRODUCTION

Cardiovascular disease acts as the leading cause of death in the world^1^. Atherosclerotic disorders, especially atherosclerosis, severely threaten people’s health with respect to its high morbidity and mortality^2^. As a complex chronic inflammatory disease, etiology of atherosclerosis initiates in endothelial inflammation and injury^3^. After the activated monocyte adhesion and penetration with more modified lipoproteins infiltrating in the injured endothelium, the accumulation of inflammatory factors promotes macrophages to M1 polarization and foam cell formation, which accelerates inflammation and cell death and eventually causes atherosclerotic plaque distablilization^4, 5^.

Pyroptosis, a manner of programmed cell death (RCD), plays a pivotal role in atherosclerosis^6^. Characterized by the plasma membrane rupture mediated by N-GSDMD, pyroptotic factors activate inflammasomes like NLRP3 and then cleave Caspase-1, which subsequently processes the precursor of IL-1β and IL-18^7^. Compared with apoptosis, which is another kind of RCD, N-GSDMD mediated plasma membrane rupture releases amounts of pro-inflammatory factors, accelerating inflammation, pyroptosis and atherosclerosis^8^.

Chemokines function as signals to induce leukocyte migration in inflammatory process^9^. According to the arrangement of cystenin in the amino terminal, chemokines have been classified into four sub-families (CC, CXC, XC, CX3C)^10^. Currently, mounting studies revealed that chemokines promote atherosclerosis^11, 12^. Jenny Jongstra-Bilen et al found that hematopoietic cell-derived CCL5 recruits monocytes and increases the abundance of intimal macrophages in 3-week lesions of Ldlr^−/−^ mice but plays a minor role in 6-week lesions, indicating that myeloid cell-derived CCL5 contributes to early lesion formation^13^. Jia-Hui Gao et al revealed that CXC chemokine ligand 12 (CXCL12) increases the levels of phosphorylated glycogen synthase kinase 3β (GSK3β) and the phosphorylation of β-catenin at the Thr120 position by interacting with CXC chemokine receptor 4 (CXCR4), which decreases the expression of TCF21 and ABCA1, inhibiting ABCA1-dependent cholesterol efflux from macrophages and then aggravating atherosclerosis^14^. As a 9KD secretory protein, HCC-1 (Hemofiltrate CC chemokine-1), also named CCL14, is an important member of CC-type chemokines^15^. Liang et al revealed that HCC-1 modulates cell cycle and promotes apoptosis in hepatocellular carcinoma (HCC) cell line^16^. Sean M Bagshaw et al found that urinary HCC-1 could predict persistent acute kidney injury (AKI)^17^. Meanwhile, Christina Massoth et al demonstrated that elevated HCC-1 levels predict persistent acute kidney injury (AKI) in cardiac surgery patients with moderate or severe acute kidney injury^18^. However, the role of HCC-1 in atherosclerosis is still unclear and has not been reported.

In this study, we aimed to investigate the role of HCC-1 in atherosclerosis. Our results firstly demonstrated that HCC-1 expression was markedly increased in serum and atherosclerotic plaque in patients with atherosclerosis. Furthermore, we indicated that HCC-1 overexpression exacerbated atherosclerosis in ApoE^−/−^ mice and promoted THP-1 monocytes adhesion, M1 polarization and induced inflammation and pyroptosis both in ECs and M1 Macrophages in vitro. Our findings revealed the promising potential of HCC-1 to serve as a early biomarker for the diagnosis and stratification of atherosclerosis.

## MATERIALS AND METHODS

The data that support the findings of this study are available from the corresponding authors upon reasonable request.

### Human tissues and serum Samples

Atherosclerotic plaque samples were collected from patients who underwent carotic endarterectomy at Nanfang Hospital (Guangdong, China). All individuals agreed to participate in the experiment and signed an informed consent. This experiment was approved by the Ethics Committee of Nanfang Hospital (Ethics approval ID: NFEC-2018-142).

Serum from 33 disease free-individuals and from 69 patients diagnosed with coronary atherosclerosis, 10 patients diagnosed with atherosclerosis and diabetes and 13 patients diagnosed with atherosclerosis and hyperlipidemia were obtained from NanFang Hospital (Guangdong, China). Exclusion criteria were as follows: patients with malignant tumors; patients with severe liver and kidney dysfunction; patients with other infectious or immune diseases; patients with communication and cognitive dysfunction.

### Enzyme-linked immunosorbent assay (ELISA)

To assess the HCC-1 levels, human serum samples were prepared and cell culture supernatants were harvested after cultured for 48h. Samples were detected by HCC-1 Human ELISA Kit (ELH-HCC1-1, Raybiotech, United States) according to the manufacturer’s protocols.

### Mice

All animal experiments were approved by the Animal Experiment Committee of Nanfang Hospital at Southern Medical University (Guangzhou, China). 6-week-old ApoE^−/−^ mice on C57BL/6J background were obtained from “GemPharmatech Co., Ltd”. Mice are all male. Adeno-associated virus serotype 9 (AAV9) vectors overexpressing HCC-1 or negative control AAV9 vector that only contains DNA sequence for Green Fluorescent Protein (GFP) (AAV9-GFP) were generated by HANBIO Technology (Shanghai, China). The mice were randomly divided into two groups (6-8 mice per group) and infected with AAV9-HCC-1 or AAV9-GFP via the tail vein. After fed with Western high-fat diet for 12 weeks, mice were sacrificed and tissues and serums were collected for further analysis. This experiment was approved by the Ethics Committee of Nanfang Hospital.

### Quantification of Atherosclerosis Burden

For mouse atherosclerosis study, mice were euthanized by CO2 inhalation and perfused with PBS via the left ventricle for 5 minutes. After perfusion, the hearts and aortas were collected and prepared for further analysis. For en face analysis, aortas were isolated with gentle removal of perivascular fat and fixed in 10% neutralized formalin for overnight. Aortas were opened longitudinally along the ventral midline from the iliac arteries to the aortic root, pinned out flat on a black surface, and stained with Oil Red O (Ab150678, Abcam, United States). The Oil Red O stained areas were traced and quantified manually using image J software. Lesion areas in aorta were shown as the ratio of plaque area/total aorta area. For aortic root lesion analysis, hearts were fixed in 10% neutralized formalin or preserved in liquid nitrogen. To determine aortic root morphometric lesion, hearts fixed in the 10% neutralized formalin were cutted for serial sections (2μm) and stained with Hematoxylin and Eosin, Sirius Red, or Masson trichrome. To determine the components of the plaque, hearts preserved in liquid nitrogen were cutted for serial cyrosections (4μm) and stained with Oil Red O, antibodies against Cleaved-caspase-1 (1:100; AF4005-50, Affinity, United States), Ly6C (1:100; E-AB-70362, Elabscience, China), P65 (1:100; AF5006, Affinity, United States), NLRP3 (1:100; DF7438, Affinity, United States) or CD68 (1:100; DF7518-50, Affinity, United States). The secondary fluorescent antibodies was Goat anti rabbit (1:1000, L114A, GeneCopoeia, United States). Nuclei were stained with DAPI (4’,6-diamidino-2-phenylindole, C1002, Beyotime, China). Stained areas were traced and quantified using image J software and data were presented as ratio of staining positive area/plaque area.

### Cell Culture and treatment

Human umbilical vein endothelial cells (HUVECs) and the human monocytic leukemia cell line, THP1, were purchased from American Type Culture Collection (ATCC). HUVECs were cultured in complete medium supplemented with 90% Dulbecco’s modified Eagle’s medium (DMEM; C11995500CP, Gibco, United States), 10% fetal bovine serum (FBS; A3160801, Gibco, United States), 100U/mL penicillin-streptomycin (15140-122, Gibco, United States). THP1 monocytes were culture in THP-1 special culture medium. All cells were incubated in humidified air containing 5% CO2 at 37°C. To simulate the function of HCC-1 in circulation in vitro, Recombinant HCC-1 protein (20ng/ml, PP-300-38B-2, Peprotech, United States) was added to cell culture medium. To simulate the influence of endothelial cells overexpressing HCC-1 on adjacent cells, co-culture system was constructed. In the co-culture system, THP-1 monocytes or THP-1 derived macrophages were cultured in the lower chamber and HUVECs were cultured at the upper chamber (0.4μm) to secrete soluble molecules HCC-1 to lower chamber. To differentiate THP-1 monocytes to resting macrophages (M0), THP-1 monocytes were treated with 20ng/ml phorbol-12-myristate-13-acetate (PMA; P8139-1MG, Sigma, United States) for 48h. M0 cells were polarized into M1 with the treatment of 20ng/ml IFN-γ (RIFNG100, Invitrogen, United States), 20ng/ml IL-6 (RP8619, Invitrogen, United States) and 10ng/ml lipopolysaccharide (LPS; tlrl-lpsbiot, Invitrogen, United States) for 48h.

### Lentivirus Construction and Infection

Lv-HCC-1, Lv-MOCK, Lv-sh-HCC-1 and Lv-sh-MOCK were constructed and synthesized by HANBIO technology (Shanghai, China). Cells were infected with Lv-HCC-1, Lv-MOCK, Lv-sh-HCC-1 and Lv-sh-MOCK at a multiplicity of infection of 50 infection units per cell in the presence of 8 μg/mL of polybrene. After 72h, cells were digested, seeded and selected in the presence of 8μg/mL of puromycin to construct the stable infected cell line.

### Cell transfection

TLR4 silencing vector (si-TLR4) and negative control vectors (Scrambled) were purchased from Ribo Targets (Guangzhou, China). HUVECs and THP-1 monocytes were transfected with 50 nM si-TLR4 or si-control vectors for 6h and then washed and cultured with fresh complete medium. qRT-PCR and Western blot (WB) were performed to observe the efficiency of siRNA knockdown.

### In Vitro Monocyte Adhesion Assay

After centrifuged and washed by PBS in two times and re-suspended by 1ml 0.1% BSA, THP-1 cells were treated with 2ul CFSE (65-0850-84, Invitrogen, United States) or Dil (D3911, Invitrogen, United States) in dark for 20 minutes. After incubation, RPMI 1640 medium (C11875500CP, Gibco, United States) supplemented with 20% fetal bovine serum was added to stop the reaction on ice. After washed by complete medium two times and calculating the cell number to 2 millions, THP-1 cells were added to HUVEC and incubated for 4h in incubator. Non-adherent cells were washed away by PBS in three times and THP-1 cells on the top of HUVEC monolayer were photographed under a fluorescence microscope (Nikon; Japan).

### Western Blot Analysis

For western blot analysis, adhered cells were lysed and whole cell, cytosolic, and nuclear protein was extracted and subsequently estimated by BCA colorimetric assay kit (23227, Pierce, United States) according to manufacturer guidelines. Equal amount of proteins (80ug) were loaded and separated by 10% and 12% SDS-PAGE gel. After electrophoresis, proteins were electro-transferred to nitrocellulose membranes and blocked by 5% BSA for 2h and subsequently incubated with primary antibodies at 4°C for 18h. After incubation with primary antibodies, membranes were washed by TBST and incubated with HRP-conjugated secondary antibody (1:5000) for 2 h. After washing with TBST, protein bands were visualized with a hypersensitive ECL chemiluminescence kit (NCM Biotech, China). Densitometry was quantified by image J software.

### Immunohistochemistry (IHC)

IHC was performed as per standard protocol reported earlier^19^. After blocked with immune serum, sections were incubated with primary abtibodies against HCC-1 (1:100; DF13362, Affinity, United States) and TNF-α(1:100; AF7014, Affinity, United States) at 4° overnight. The following procedure was performed with the PolyExcel HRP/DAB Detection System kit (PathnSitu Biotechnologies) according to the manufacturer instructions. Immune reactions were visualized by adding the DAB (3,3′-diaminobenzidine tetrachloride) and all sections were counterstained with hematoxylin. Relative protein expression was quantified by image J software.

### Quantitative Real-Time Polymerase Chain Reaction

For total RNA extraction, cells were lysed by TRIzol reagent (15596018, Invitrogen, United States) according to the manufacturer guidelines and cDNA was synthesized using Reverse Transcription Kit (R222-01, Vazyme, China). Quantitative real time PCR was performed on LightCycler 480 Instrument II (Roche, Pleasanton, CA, USA) using SYBR Green Polymerase Chain Reaction kit (Q131-02, Vazyme, China) and all reactions were complete in triplicate. The mRNA expression of individual genes were normalized to the expression of GAPDH.

### DNA Fragmentation Evaluation

DNA fragmentation evaluation was measured by TUNEL staining assay according to standard protocol. In brief, endothelial and M1 cells were cultured on coverslips in a 24-well plate. After treatment, cells were fixed with 4% paraformaldehyde for 30 min and then permeabilized with 0.3% Triton-X-100 for 5min in room temperature. After washed two times with PBS, cells were stained with TUNEL reaction mix (E-CK-A325, Elabscience, China) for 1h at 37°C in dark and then stained by DAPI for 5min. Cells were photographed under a fluorescence microscope (Nikon; Japan).

### LDH release assay

The supernatants from treated cells were centrifuged and collected for LDH release assay following the manufacturer’s instructions (AC10187, ACMEC, China). The absorbance was measured at 450 nm on a spectrophotometric microplate reader.

### Cell Death Assay

Hoechst 33352/PI staining assay was used to assess the pyroptotic cell death in HUVEC and M1 cells. For hoechst 33342/PI staining, cell stain buffer and the equal amount of ddH20 was mixed to prepare the working fluid. After washed with PBS, cells were incubated with Hoechst 33352/PI working fluid (PH0532, Phgene, China) at 4°C on ice for 30min. After incubation, cells were washed and then photographed under a fluorescence microscope (Nikon; Japan).

### Statistical Analysis

Data were analyzed using GraphPad Prism 8.0 software and presented as mean±SD. Each experiment was repeated at least three times. Shapiro-Wilks test was used to test normality of data distribution. Unpaired t-test was used for normally distributed data and Mann-Whitney U test was used for non-normally distributed data. For multiple groups testing, ANOVA analysis was conducted as 1-way ANOVA depending on normal distribution with Tukey post hoc test or Kruskal-Wallis test depending on non-normal distribution with Dunn’s multiple comparisons for correction. In all statistical analyses, P values below 0.05 were considered to indicate a statistically significant result.

## RESULTS

### HCC-1 expression increased in human atherosclerotic plaque

To assess HCC-1 expression in patients with atherosclerosis, human atherosclerotic plaques and normal arterial tissues were randomly collected. In IHC analysis, the results showed that HCC-1 expression was higher in atherosclerotic plaque versus control (Figure 1A and 1B). Three normal arterial tissues and six atherosclerotic plaques from AS patients were randomly chosen to examine HCC-1 mRNA expression. RT-PCR analysis showed that the mRNA expression levels of HCC-1 were remarkably enhanced by 17.6-fold in atherosclerotic plaques in comparison with normal arterial tissues (Figure 1C). To evaluate the protein expression of HCC-1 in atherosclerotic plaques, western blot analysis showed that there was a 4.53-fold increase in HCC-1 protein levels in atherosclerotic plaques compared with control (Figure 1D and 1E). These findings suggested that human atherosclerotic plaque had a marked increase in HCC-1 both in protein and mRNA levels.

**Figure 1.**
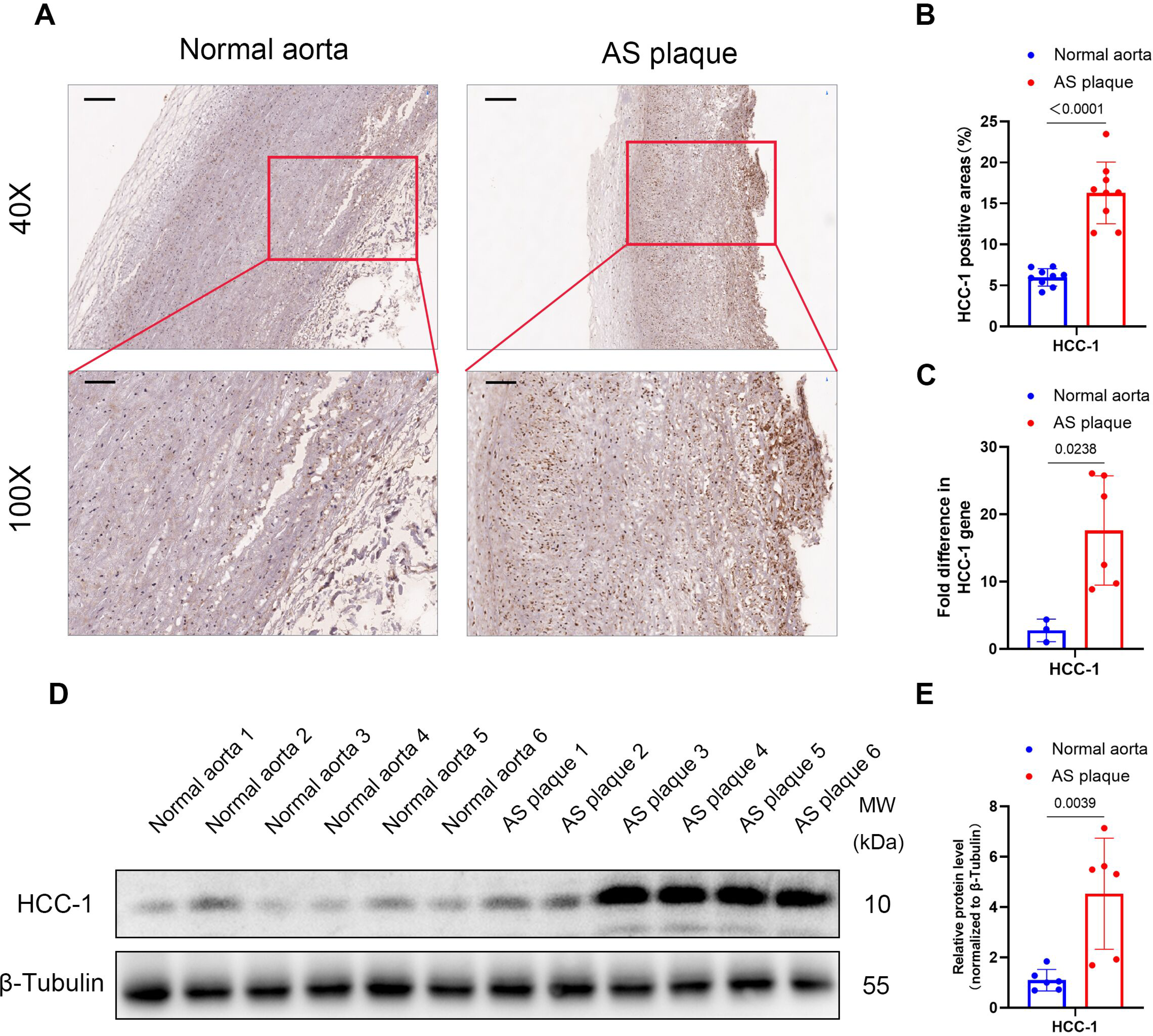
HCC-1 (Hemofiltrate CC chemokine-1) expression increased in human atherosclerotic lesions. Human atherosclerotic plaques and normal arterial tissues were randomly collected. **A** and **B**, Representative images of IHC analysis (**A**) and quantification (**B**) of HCC-1 in Human atherosclerotic plaques and normal arterial tissues (n=9 per group, unpaired t test, P<0.0001, Shapiro-Wilks test [SW] showed normal distribution). **C**, mRNA expression levels of HCC-1 in normal arterial tissues and Human atherosclerotic plaques (normal n=3 vs AS n=6, Mann-Whitney test, P=0.0238, SW showed no normal distribution) and GAPDH was used as an internal loading control. **D** and **E**, protein expression levels (**D**) and quantification (**E**) of HCC-1 in normal arterial tissues and Human atherosclerotic plaques (n=6 per group, unpaired t test, P=0.0039, SW showed normal distribution) and β-Tubulin was used as an internal loading control. Data were exhibited as mean ± SD. Scale bar=250um for (40X) and 100um for (100X).

### HCC-1 exacerbated atherosclerotic lesion formation in ApoE^−/−^ mice

To determine the role of HCC-1 in the progression of atherosclerosis in vivo, ApoE^−/−^ mice were treated with AAV9-Control and AAV9-HCC-1 under a high fat diet for 12 weeks and their atherosclerotic lesions and serum samples were evaluated. The results of en face analysis showed that ApoE^−/−^ mice treated with AAV9-HCC-1 had a higher average lesion areas than controls (Fig. 2A). In aortic root section analyses, the results revealed that HCC-1 overexpression increased atherosclerotic lesions area (Fig. 2B and 2C), lipid deposition (Fig. 2D and 2E), whereas decreased collagen contents in ApoE^−/−^ mice (Fig. 2F through 2I). These results suggested that HCC-1 exacerbated atherosclerosis in ApoE^−/−^ mice.

**Figure 2.**
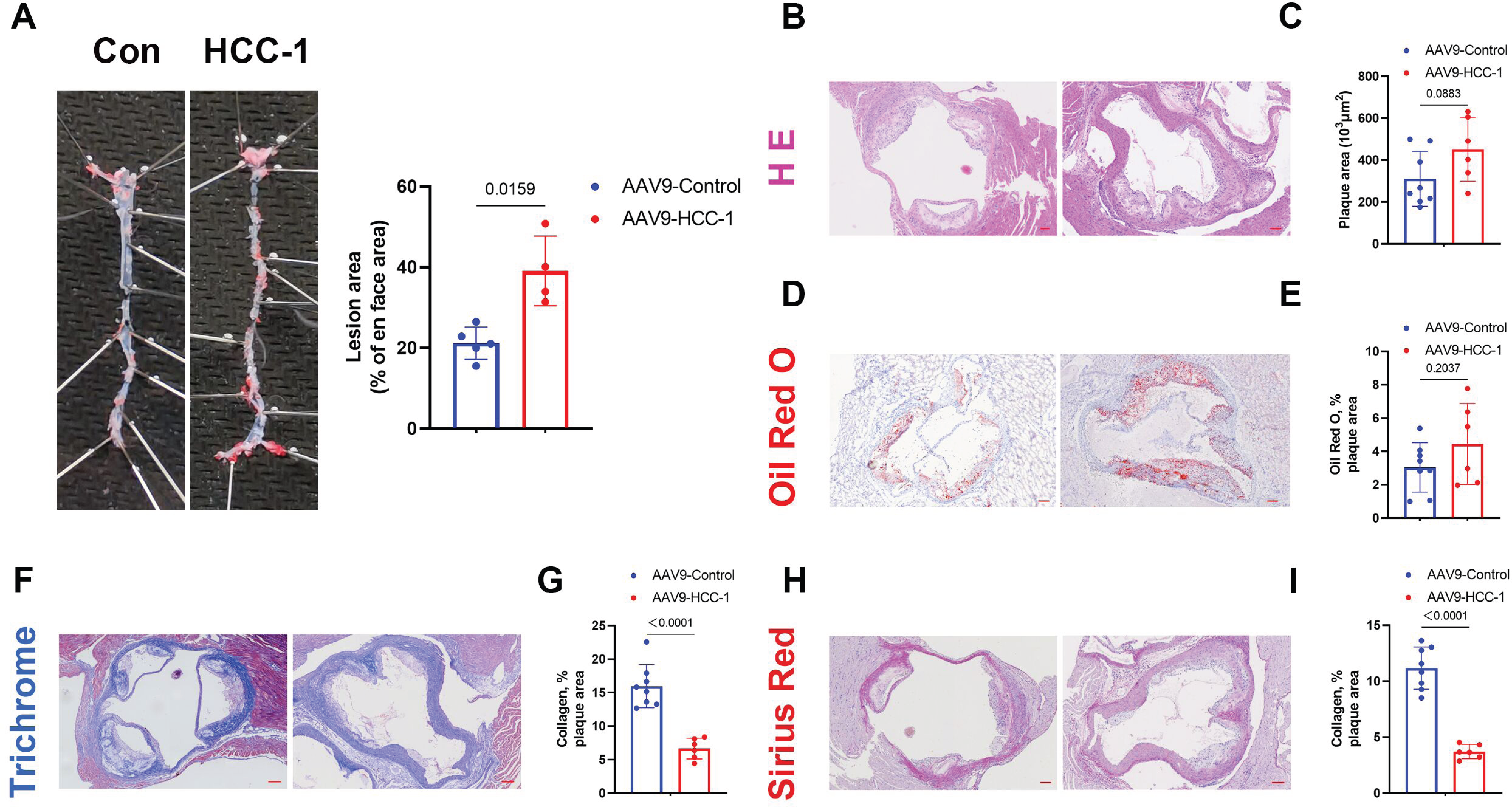
HCC-1 (Hemofiltrate CC chemokine-1) exacerbated atherosclerotic lesion formation in ApoE^−/−^ mice. ApoE^−/−^ mice treated with adeociated-virus-9 (AAV9)-mediated overexpression of HCC-1 (AAV9-HCC-1 mice) and the controls (AAV9-Control mice) were fed with high fat diet for 12 weeks. **A**, Oil red O staining of aorta en face in different groups (Control n=5 vs AAV9-HCC-1 n=4, Mann-Whitney test, P=0.0159, Shapiro-Wilks [SW] showed no normal distribution). **B**-**I**, H&E (**B**), Oil red O (**D**), Masson trichrome (**F**) and sirius red (**H**) staining of aortic root cross sections and quantification of lesion areas (**C**), Oil red O-positive lesion areas (**E**) and plaque collagen content (**G** and **I**) (Control n=8 vs AAV9-HCC-1 n=6, unpaired t test, SW showed normal distribution). Data were exhibited as mean ± SD. Scale bar=100μm.

### Pro-inflammatory stimulus increased HCC-1 expression in HUVECs and THP-1 monocytes

Knowing that HCC-1 expression was up-regulated in atherosclerotic lesions, we next studied the expression of HCC-1 in the condition of inflammation or hyperlipidemia because inflammatory factors, like LPS and Ox-LDL, play a critical role in atherosclerosis. ELISA analysis revealed that HCC-1 secretion was enhanced in HUVECs treated by LPS (Figure 3A), while in THP-1 monocytes, Ox-LDL was the main factor to promote the secretion of HCC-1 (Figure 3B). To study the gene expression, RT-PCR was performed and the results indicated that both LPS and Ox-LDL up-regulated HCC-1 gene expression in HUVECs and THP-1 monocytes (Figure 3C and 3D). In order to simulate the real condition, Co-culture system was established to study the influence of HUVECs overexpressing HCC-1 on THP-1 monocytes, M0 macrophages and M1 macrophages. The results suggested that HCC-1 gene and protein expression were up-regulated in THP-1 monocytes, M0 macrophage and M1 macrophage after cocultured with HUVECs overexpressing HCC-1 (Figure 3E through 3H), revealing that ECs, which was overexpressing HCC-1 in the atherosclerotic condition, could secrete inflammatory factors including HCC-1 to interact with THP-1 monocyte, M0 macrophage and M1 macrophage and then enhance their HCC-1 expression.

**Figure 3.**
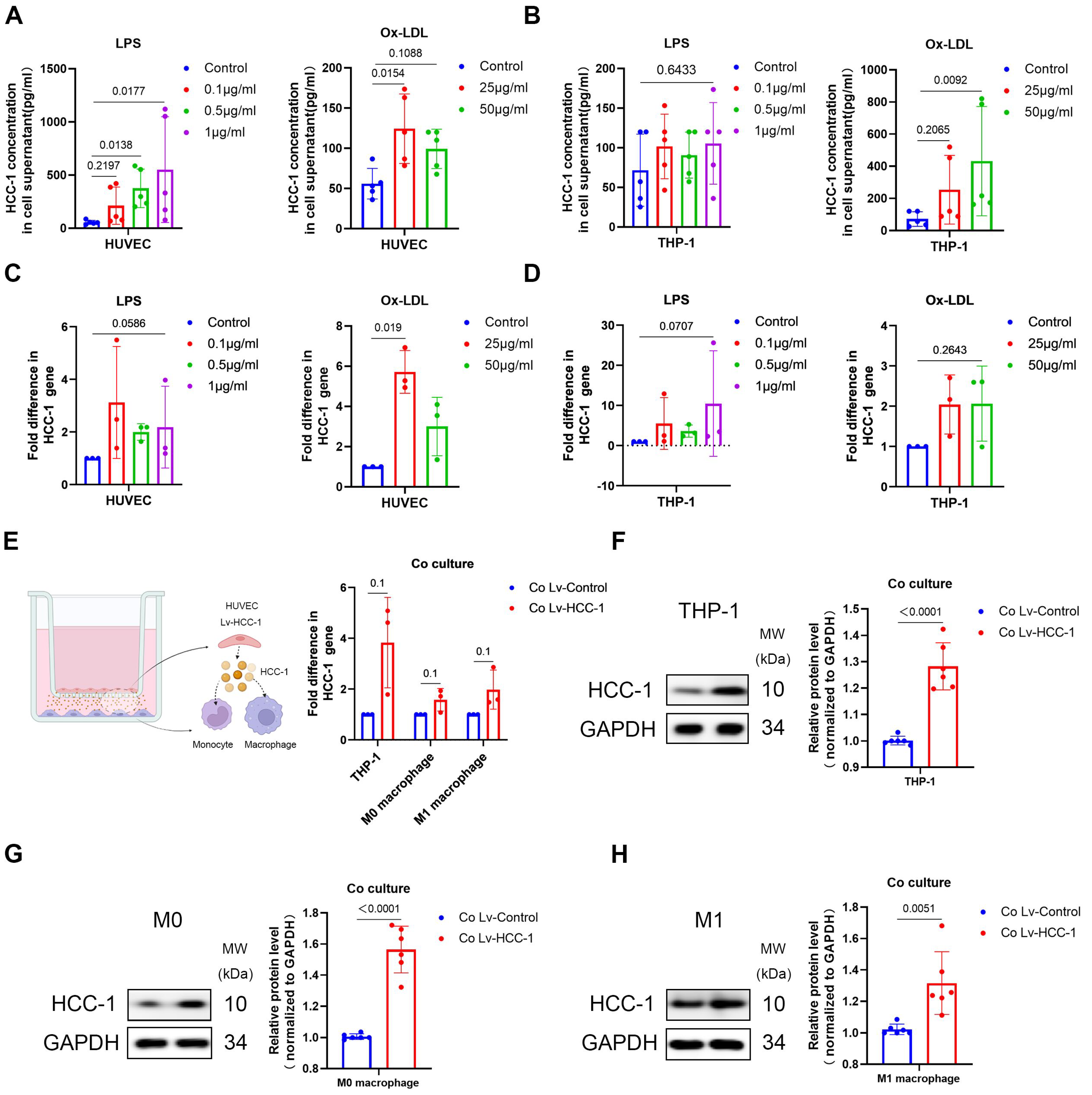
Pro-inflammatory stimulus increased HCC-1 (Hemofiltrate CC chemokine-1) expression in HUVECs and THP-1 monocytes. HUVECs and THP-1 monocytes were pretreated with LPS (0.1, 0.5, 1μg/ml) or Ox-LDL (25, 50μg/ml) for 48h. THP-1 monocytes, THP-1 derived M0 Macrophages and THP-1 derived M1 Macrophages were cocultured with HUVECs-Lv-HCC-1 for 48h. **A** and **B**, HCC-1 concentration in supernatant of cultured HUVECs (**A**) and THP-1 monocytes (**B**) after treated with LPS and Ox-LDL (n=5 per group, Kruskal-Wallis test with Dunn’s multiple comparison test, Shapiro-Wilks [SW] did not show normal distribution). **C** and **D**, HCC-1 gene expression of cultured HUVECs (**C**) and THP-1 monocytes (**D**) after treated with LPS and Ox-LDL (n=3 per group, Kruskal-Wallis test with Dunn’s multiple comparison test, P=0.0036 in Ox-LDL treated HUVEC group, SW did not show normal distribution). **E**, Diagram of the co-culture system and HCC-1 gene expression of THP-1 monocytes, M0 macrophages and M1 macrophages after cocultured with HUVECs overexpressing HCC-1 (n=3 per group, Mann-Whitney test, P=0.1, SW did not show normal distribution). **F**, **G** and **H**, Expression of HCC-1 in THP-1 monocytes (**F**), M0 macrophages (**G**) and M1 macrophages (**H**) after cocultured with HUVECs overexpressing HCC-1 was quantitatively analyzed, and GAPDH was used as an internal loading control (n=6 per group, unpaired t test, SW showed normal distribution). Data were exhibited as mean ± SD.

### HCC-1 promoted THP-1 monocytes to adhere to endothelial cells

To investigate how HCC-1 contributes to atherosclerosis, lentivirus encoding HCC-1 was constructed and recombinant HCC-1 protein and HUVECs-Lv-HCC-1 co-culture system were used to simulate the function of exogenuous HCC-1. ELISA analysis showed that HCC-1 secretion in HUVECs and THP-1 monocytes was greatly enhanced by lentivirus encoding HCC-1 (Figure SI). Leukocyte adhesion plays a pivotal role in the progression of atherosclerosis^20^. To determine whether HCC-1 participated in the process of monocytes adhesion, adhesion experiment was performed and the results showed that treatment of recombinant HCC-1 protein in HUVECs or THP-1 monocytes both promoted THP-1 monocytes to adhere to endothelial cells (Figure 4A and 4B). It is reported that adhesion molecules, like ICAM-1, VCAM-1 or P-selectin expressed by endothelial cells, or metallopeptidases (MMPs) expressed by THP-1 monocytes, exerted critical function in monocytes adhesion^21–23^. RT-PCR and Western blot analysis showed that recombinant HCC-1 protein up-regulated the protein expression of P-selectin, ICAM-1 and VCAM-1 in HUVECs and MMP-7 and MMP-10 in THP-1 monocytes (Figure 4C and 4D, Figure SI), suggesting that HCC-1 protein was implicated in the process of monocytes adhesion. Meanwhile, we observed similar results by overexpressing HCC-1 via lentivirus encoding HCC-1 both in HUVECs and THP-1 monocytes (Figure 4E through 4I, Figure SI)

**Figure 4.**
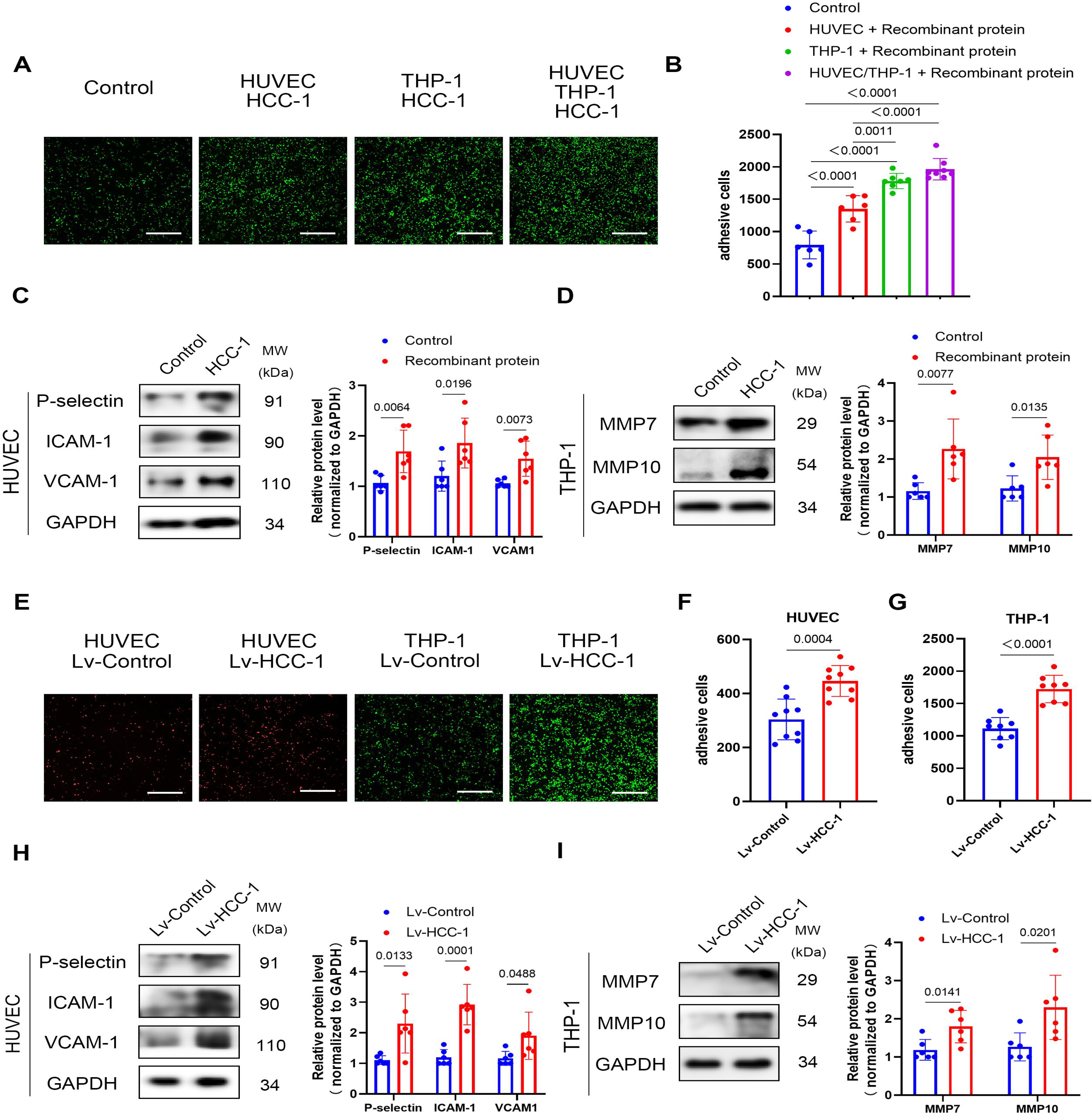
HCC-1 (Hemofiltrate CC chemokine-1) promoted THP-1 monocytes to adhere to endothelial cells. HUVECs and THP-1 monocytes were pretreated with recombinant HCC-1 protein (20ng/ml) for 48h or infected with lentivirus encoding HCC-1. **A**, Representative images of adhered THP-1 monocytes stained with carboxifluorescein diacetate succinimidyl ester (CFSE). **B**, Quantification of adhered THP-1 monocytes (Statistical significances analyzed by 1-way ANOVA with Tukey post hoc test, P<0.0001, Shapiro-Wilks [SW] showed normal distribution). **C** and **D**, Expression of cell adhesion proteins (**C**) and matrix metallopeptidase (MMP) (**D**) was quantitatively analyzed, and GAPDH was used as an internal loading control (n=6 per group, unpaired t test, SW showed normal distribution) (the internal loading control in **C** is shared in Fig.6A and Fig.7A as the same batch of result). **E**, Representative images of adhered THP-1 monocytes stained with carboxifluorescein diacetate succinimidyl ester (CFSE) or Dil. **F** and **G**, Quantification of adhered THP-1 monocytes in HUVECs-Lv-HCC-1 group (**F**) (n=9 per group, unpaired t test, P=0.0004, SW showed normal distribution) or THP-1-Lv-HCC-1 group (**G**) (n=8 per group, unpaired t test, P<0.0001, SW showed normal distribution). **H** and **I**, Expression of cell adhesion proteins (**H**) and matrix metallopeptidase (MMP) (**I**) was quantitatively analyzed, and GAPDH was used as an internal loading control (n=6 per group, unpaired t test, SW showed normal distribution) (the internal loading control in **H** is shared in Fig.**S3C** and Fig.**S6G** as the same batch of result). Data were exhibited as mean ± SD. Scale bar=1mm.

### HCC-1 promoted macrophages to M1 polarization

Na Li et al reported that HCC-1 had a function in inhibiting colon cancer cell by suppressing M2 polarization of tumor-associated macrophage^24^. But in their study, they used 1μg/ml HCC-1 to treat M0 macrophages, which is exceedingly higher than normal working concentration (20ng/ml). Therefore, their consequence might be inaccurate and we assessed the function of HCC-1 in M1 polarization of macrophages again. To investigate whether HCC-1 promoted M0 macrophages to M1 polarization so as to accelerate atherosclerosis, con-focal microscope was used to assess the co-localization of HCC-1 and M1 macrophage in human atherosclerotic plaque. The results showed that HCC-1 expressed by macrophages colocalized with M1 marker including TNF-α, CD86, IL-1β, IL-6 and NOS2, indicating that HCC-1 could promote M0 macrophages to M1 polarization in human atherosclerotic plaque (Figure 5A, Figure SII). Meanwhile, Western blot analyses showed that HUVECs-Lv-HCC-1 or recombinant HCC-1 protein had a capacity to enhance the expression of M1 marker and decrease the expression of M2 marker in M0 macrophage, indicating that HCC-1 promoted M0 macrophages to M1 polarization (Figure 5B through 5E).

**Figure 5.**
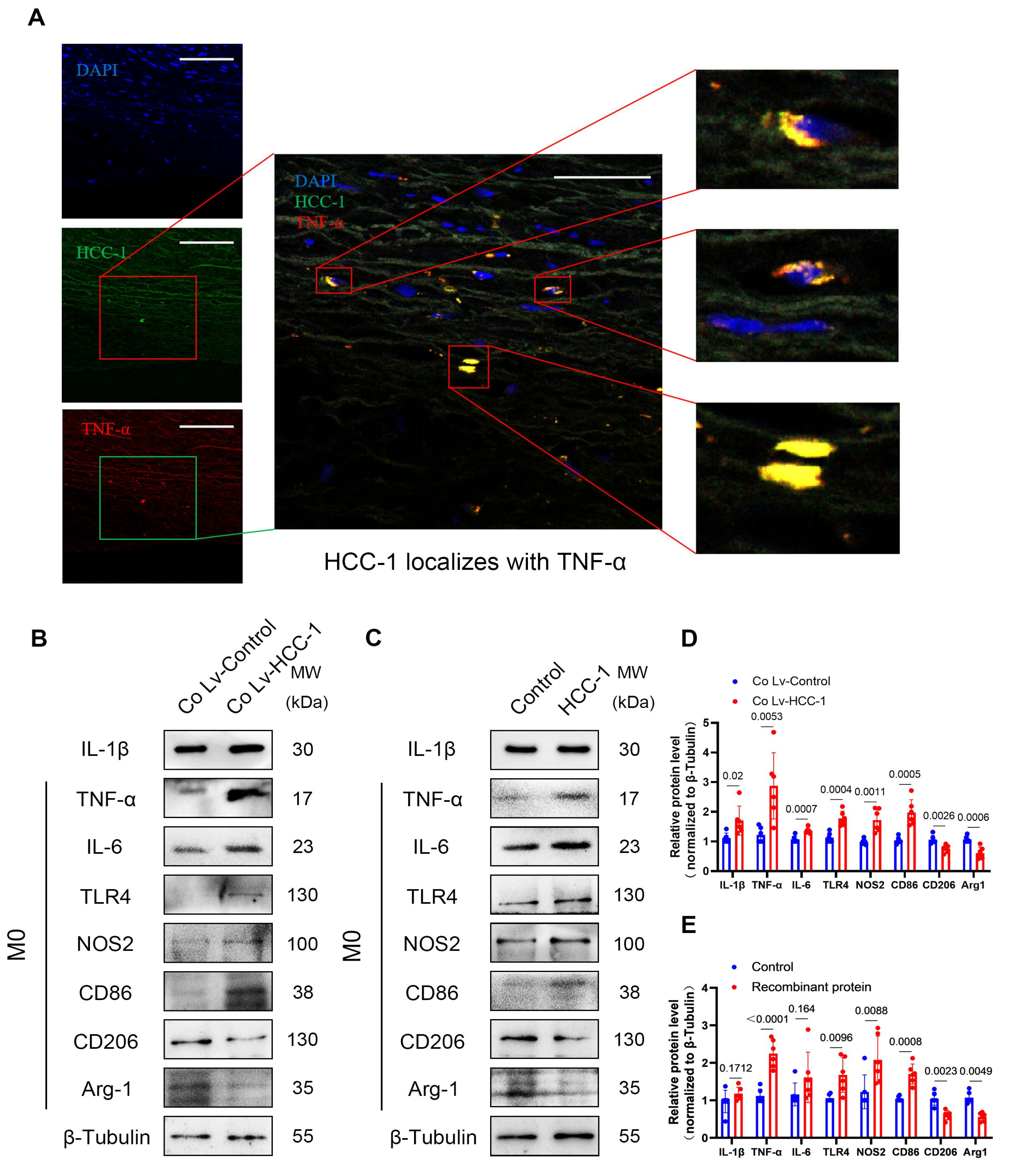
HCC-1 (Hemofiltrate CC chemokine-1) promoted macrophages to M1 polarization. **A**, Representative image of co-localization between HCC-1 and TNF-α under Con-focal microscope. **B** - **E**, Protein expression of IL-1β, TNF-α, IL6, TLR4, NOS2, CD86, CD206, Arg1 in M0 macrophages co-cultured with HUVECs overexpressing HCC-1 (**B**) or treated with recombinant HCC-1 protein (20ng/ml) (**C**) were quantitatively analyzed, and β-Tubulin was used as an internal loading control (**D** and **E**) (n=6 per group, unpaired t test, Shapiro-Wilks [SW] showed normal distribution). Data were exhibited as mean ± SD. Scale bar=100um for (left) and 50u for (middle).

### HCC-1 induced inflammation and pyroptosis in endothelial cells and M1 Macrophages

Knowing that HCC-1 could promote monocytes adhesion and M1 polarization, we hypothesized that HCC-1 might promote endothelial cells or Macrophages inflammation by inducing cell injury since injured endothelial cell and macrophages secreted amounts of pro-inflammatory factors and formed an inflammatory cascade, resulting in further recruitment of monocytes, stronger inflammation and accelerated plaque progression^5, 25^. Of note, as a kind of pro-inflammatory programmed cell death, pyroptosis leads to marked release of inflammatory factors and play a pivotal role in atherosclerosis^8^. To confirm our hypothesis, Western blot and RT-PCR analyses were performed to assess the expression levels of inflammation and pyroptosis related molecules in HUVECs and M1 Macrophages after treated with recombinant HCC-1 protein or infected by lentivirus encoding HCC-1 or cocultured with HUVECs-Lv-HCC-1. The results showed that both recombinant HCC-1 protein, lentivirus encoding HCC-1 and HUVECs-Lv-HCC-1 enhanced the expression of inflammatory factors like IL-6 and TNF-α, and the expression of pyroptosis associated proteins like cleaved-IL-1β、cleaved-caspase-1、ASC and GSDMD-N-terminal in HUVECs and M1 Macrophages (Fig. 6A, 6B, 6E and 6F, Figure SIII, SIV and SV) and lentivirus encoding sh-HCC-1 decreased the expression of inflammatory factors (Figure SV). To further dissect pyroptosis in ECs and M1 Macrophages, LDH release assay and PI and TUNEL staining were respectively evaluated. The results showed that PI and TUNEL positive rate were remarkably enhanced in HUVECs and M1 Macrophages after treated with recombinant HCC-1 protein, lentivirus encoding HCC-1 or cocultured with HUVEC-Lv-HCC-1 (Fig. 6C, 6D, G and 6H, Figure SIII, SIV and SV). Interestingly, LDH release was enhanced in HUVECs infected with lentivirus encoding HCC-1 and M1 Macrophages treated with recombinant HCC-1 protein or infected with lentivirus encoding HCC-1, but not in HUVECs treated with recombinant HCC-1 protein. These results showed that HCC-1 could induce inflammation and pyroptosis in endothelial cells and M1 Macrophages.

**Figure 6.**
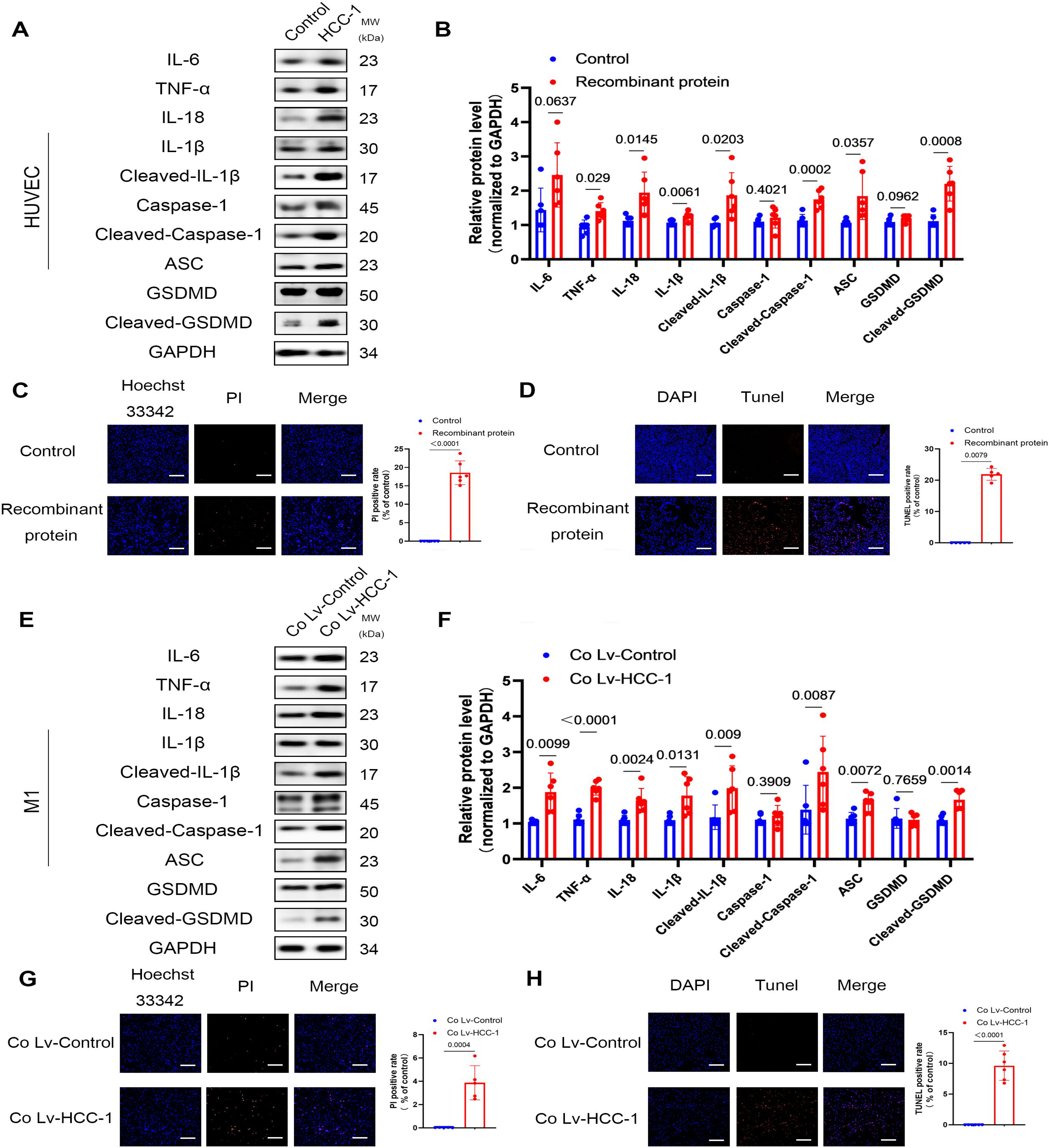
HCC-1 (Hemofiltrate CC chemokine-1) induced inflammation and pyroptosis in endothelial cells and M1 Macrophages. HUVECs were pretreated with recombinant HCC-1 protein (20ng/ml) for 48h or infected with lentivirus encoding HCC-1. M1 Macrophages were pretreated with recombinant HCC-1 protein (20ng/ml) for 48h or infected with lentivirus encoding HCC-1 or cocultured with HUVECs-Lv-HCC-1 for 48h. **A** and **E**, Protein expression levels of IL-6, TNF-α, IL-18, IL-1β, Cleaved-IL-1β, caspase-1, Cleaved-caspase-1, ASC, GSDMD and Cleaved-GSDMD in HUVECs treated with recombinant HCC-1 protein (**A**) (the internal loading control in **A** is shared in Fig.4C and Fig.7A as the same batch of result) or in M1 Macrophages cocultured with HUVECs-Lv-HCC-1 (**E**) (the internal loading control in **E** is shared in Fig.7E as the same batch of result) were determined. **B** and **F**, Expression of inflammatory and pyroptosis-related proteins in HUVECs treated with recombinant HCC-1 protein (**B**) or in M1 Macrophages cocultured with HUVECs-Lv-HCC-1 (**F**) were quantitatively analyzed, and GAPDH was used as an internal loading control (n=6 per group, unpaired t test, Shapiro-Wilks [SW] showed normal distribution). **C** and **G**, Representative images of Hoechst33342/PI double staining assay and the quantification of PI positive rate (normalized to average PI positive rate in control group) in HUVECs treated with recombinant HCC-1 protein (**C**) (n=6 per group, unpaired t test, P<0.0001, SW showed normal distribution) or in M1 Macrophages cocultured with HUVECs-Lv-HCC-1 (**G**) (n=5 per group, unpaired t test, P=0.0004, SW showed normal distribution). **D** and **H,** Representative images of TUNEL staining assay and the quantification of TUNEL positive rate (normalized to average TUNEL positive rate in control group) in HUVECs treated with recombinant HCC-1 protein (**D**) (n=5 per group, Mann-Whitney test, P=0.0079, SW showed no normal distribution) or in M1 Macrophages cocultured with HUVECs-Lv-HCC-1 (H) (n=6 per group, unpaired t test, P<0.0001, SW showed normal distribution). Data were exhibited as mean ± SD. Scale bar=1mm.

### HCC-1 induced inflammation and pyroptosis in endothelial cells and M1 Macrophages through NF-κB/NLRP3/Caspase-1 pathway

To investigate the underlying mechanism that how HCC-1 induced inflammation and pyroptosis in HUVECs and M1 macrophages,our findings revealed that in HUVECs, both recombinant HCC-1 protein and lentivirus encoding HCC-1 enhanced the protein and gene expression of TLR4, NLRP3 and promoted NF-κB nuclear translocation (Fig. 7A and 7B, Figure SVI) and lentivirus encoding sh-HCC-1 decreased the protein expression of TLR4, NLRP3 and inhibited NF-κB nuclear translocation (Figure SVII). To confirm whether HCC-1 induced inflammation and pyroptosis in HUVECs via TLR4/NF-κB/NLRP3/Caspase-1 pathway, we knocked down TLR4 in HUVECs by performing si-TLR4 transfection. The results of Western blot analysis showed that the expression of NLRP3, cleaved-caspase-1 and GSDMD-N-terminal was down-regulated after si-TLR4 transfection (Fig. 7C and 7D, Figure SVI), indicating that HCC-1 accurately induced inflammation and pyroptosis in endothelial cells via TLR4/NF-κB/NLRP3/Caspase-1 pathway. In M1 macrophages, Western blot and RT-PCR analyses showed that treatment of recombinant HCC-1 protein, coculture with HUVECs-Lv-HCC-1 and infection of lentivirus encoding HCC-1 enhanced the protein expression of NLRP3 and promoted NF-κB nuclear translocationin M1 Macrophages (Fig. 7F, Figure SVII), while merely HUVECs-Lv-HCC-1 enhanced the protein expression of TLR4 in M1 Macrophages (Fig. 7E), indicating that HCC-1 induced M1 macrophages pyroptosis through NF-κB/NLRP3/Caspase-1 pathway.

**Figure 7.**

HCC-1 induced inflammation and pyroptosis in endothelial cells and M1 Macrophages through NF-κB/NLRP3/Caspase-1 pathway. HUVECs were pretreated with recombinant HCC-1 protein (20ng/ml) or infected with lentivirus encoding. THP-1 derived M1 Macrophages were pretreated with recombinant HCC-1 protein (20ng/ml) or infected with lentivirus encoding HCC-1 or cocultured with HUVECs-Lv-HCC-1 for 48h. **A**, Protein expression levels of TLR4 and NLRP3 in HUVECs treated with recombinant HCC-1 protein (20ng/ml) were determined and quantitatively analyzed, and GAPDH was used as an internal loading control (n=6 per group, unpaired t test, Shapiro-Wilks [SW] showed normal distribution) (the internal loading control in **A** is shared in Fig.4C and Fig.6A as the same batch of result). HUVECs were seeded for 12h. After 30min of recombinant HCC-1 protein (20ng/ml) incubation, both cytosolic and nuclear proteins were isolated. **B**, Protein expression levels of P65 and p-P65 in HUVECs treated with recombinant HCC-1 protein (20ng/ml) were determined and quantitatively analyzed, and GAPDH and histone-H3 were used as internal loading control to cytosolic and nuclear respectively (n=6 per group, unpaired t test, SW showed normal distribution). After seeded for 12h, HUVECs were transfected with si-NC, si-TLR4 and si-TLR4 with treatment of recombinant HCC-1 protein (20ng/ml) for 48h. **C** and **D**, Protein expression levels of TLR4, NLRP3, caspase-1, Cleaved-caspase-1, GSDMD, Cleaved-GSDMD were determined and quantitatively analyzed, and GAPDH was used as an internal loading control (n=3 per group, Kruskal-Wallis test with Dunn’s multiple comparison test, P=0.0023, SW showed no normal distribution). **E**, Protein expression levels of TLR4 and NLRP3 in THP-1 derived M1 Macrophages treated with recombinant HCC-1 protein (20ng/ml) were determined and quantitatively analyzed, and GAPDH was used as an internal loading control (n=6 per group, unpaired t test, SW showed normal distribution) (the internal loading control in **E** is shared in Fig.6E as the same batch of result). THP-1 derived M0 Macrophages were induced to M1 polarization for 48h. After washed, and incubated with recombinant HCC-1 protein (20ng/ml) or cocultured with HUVECs-Lv-HCC-1 for 30min, both cytosolic and nuclear proteins were isolated. **F**, Protein expression levels of P65 and p-P65 in THP-1 derived M1 Macrophages treated with recombinant HCC-1 protein (20ng/ml) were determined and quantitatively analyzed, and GAPDH and histone-H3 were used as internal loading control to cytosolic and nuclear respectively (n=6 per group, unpaired t test, SW showed normal distribution). Data were exhibited as mean ± SD.

### HCC-1 induced inflammation and atherosclerotic plaque pyroptosis in ApoE^−/−^ mice

Because HCC-1 induced inflammation and pyroptosis in vitro, we further investigated in mice and immunofluorescence analyses showed that HCC-1 enhanced monocytes (Fig. 8A and 8B) and macrophages accumulation in plaques (Fig. 8C and 8D). Knowing that HCC-1 induced pyroptosis in vitro via NF-κB/NLRP3/Caspase-1 pathway, our results further confirmed that HCC-1 increased TUNEL positive area via activating caspase-1 (Fig. 8E through 8H), following the activation of NF-κB and NLRP3(Fig. 8I through 8L). To evaluate the level of inflammatory factors in mice, the concentration of TNF-α and IL-1β in serum were evaluated and the results revealed that HCC-1 overexpression increased the serum level of TNF-α and IL-1β (Fig. 8M and Fig. 8N) in mice. These results suggested that HCC-1 overexpression induced inflammation, pyroptosis and exacerbated atherosclerosis.

**Figure 8.**
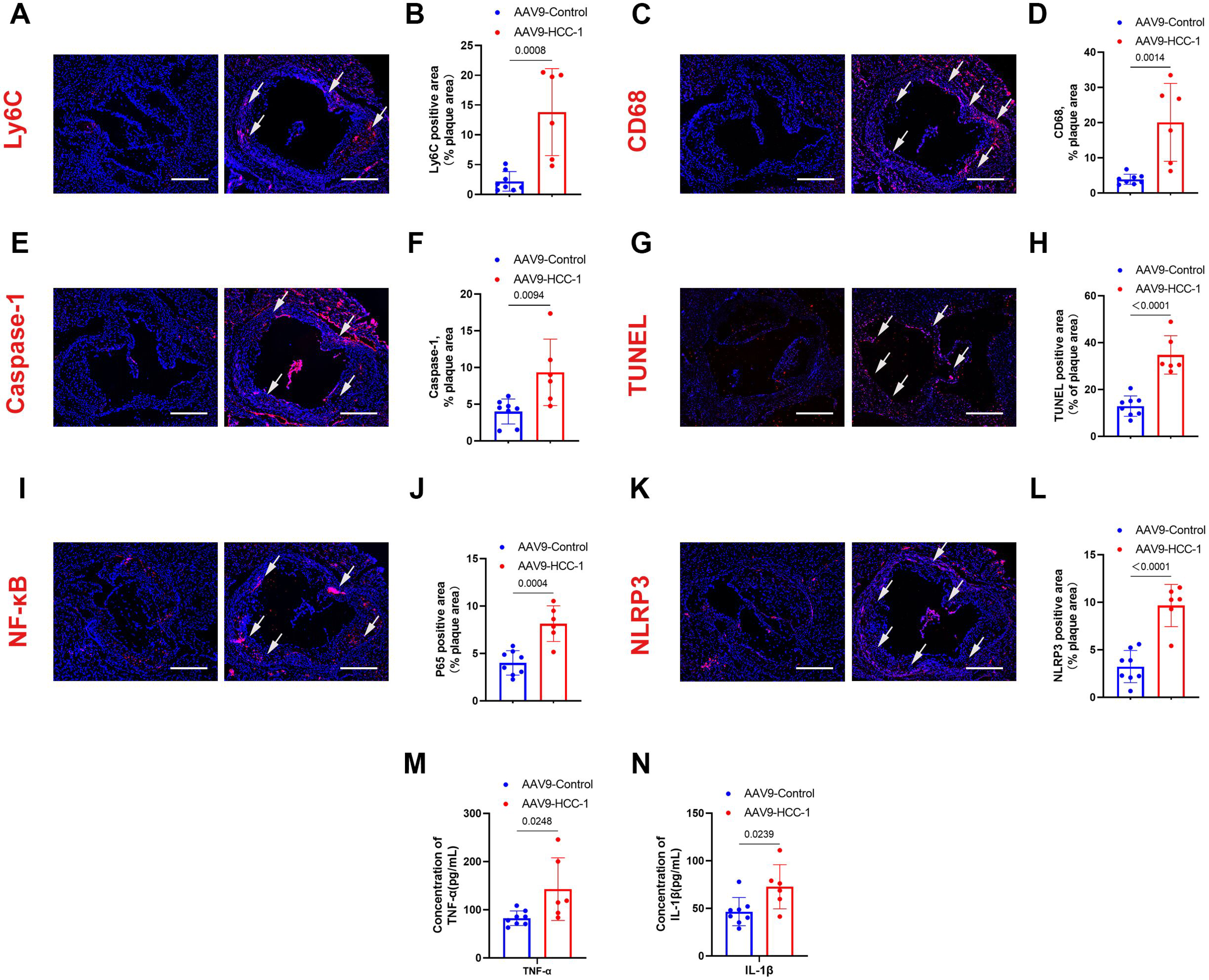
HCC-1 induced inflammation and atherosclerotic plaque pyroptosis in ApoE^−/−^ mice. ApoE^−/−^ mice treated with adeociated-virus-9(AAV9)-mediated overexpression of HCC-1 (AAV9-HCC-1 mice) and the controls (AAV9-Control mice) were fed with high fat diet for 12 weeks. **A** - **D**, Anti-LyC6 (**A**) and Anti-CD68 (**C**) antibody stained aortic root cross sections and quantification of monocytes (**B**) and macrophages (**D**) accumulation (Control n=8 vs AAV9-HCC-1 n=6, unpaired t test, Shapiro-Wilks [SW] showed normal distribution). **E** - **H**, Anti-caspase-1 antibody (**E**) and TUNEL (**G**) stained aortic root cross sections and quantification of caspase-1-positive (**F**) and TUNEL-positive rate (**H**) (Control n=8 vs AAV9-HCC-1 n=6, unpaired t test, SW showed normal distribution). **I** - **L**, Anti-NF-κB (**I**) and Anti-NLRP3 (**J**) antibody stained aortic root cross sections and quantification of NF-κB (**J**) and NLRP3-positive rate (**L**) (Control n=8 vs AAV9-HCC-1 n=6, unpaired t test, SW showed normal distribution). **M** and **N**, serum levels of TNF-α (**M**) and IL-1β (**N**) in mice (Control n=8 vs AAV9-HCC-1 n=6, unpaired t test, SW showed normal distribution). Scale bar=100μm.

### Serum Levels of HCC-1 increased in Atherosclerosis Subjects

To assess the potential of HCC-1 to serve as a biomarker of atherosclerosis, serum from control and atherosclerotic subjects had been collected and tested by HCC-1 ELISA kit. The results showed that HCC-1 concentration was markedly increased in AS subjects (Fig. 9A) and with a high diagnostic efficiency (Fig. 9B). Further studies revealed that serum levels of HCC-1 increased in patients with mild stenosis (Fig. 9C) and increased with the progression of stenosis (Fig. 9D), revealing that HCC-1 might server as a early diagnostic biomarker of atherosclerosis and might reflect the degree of stenosis. Because most patients with atherosclerosis have hyperlipidemia or diabetes, we assessed whether hyperlipidemia or diabetes influenced the concentration of HCC-1 and our results suggested that diabetes and hyperlipidemia, two main interference factors in atherosclerosis, had little impact on the concentration of HCC-1 in serum (Fig. 9E and 9F).

**Figure 9.**

Serum levels of HCC-1 (Hemofiltrate CC chemokine-1) increased in Atherosclerosis subjects. Serums from patients with atherosclerosis, atherosclerosis combined with diabetes and atherosclerosis combined with hyperlipidemia were collected and measured. **A**, Serum levels of HCC-1 in control group and atherosclerosis subjects (Control n=33 vs AS n=92, Mann-Whitney test, P<0.0001, Shapiro-Wilks [SW] showed no normal distribution). **B**, ROC curve of serum levels of HCC-1 in control group (n=33) and atherosclerosis subjects (n=92). **C**, Serum levels of HCC-1 in control group and atherosclerosis with stenosis area ≤ 50% subjects (Control n=33 vs AS with stenosis n=22, unpaired t test, P<0.0001, SW showed normal distribution). **D**, Serum levels of HCC-1 in atherosclerosis with stenosis area ≤ 50% subjects, atherosclerosis with stenosis area ≤ 51%∼75% subjects and atherosclerosis with stenosis area ≤ 76%∼100% subjects (AS with stenosis area ≤ 50% n=22 vs AS with stenosis area ≤ 51%∼75% n=13 vs AS with stenosis area ≤ 76%∼100% n=14, statistical significances analyzed by 1-way ANOVA, P=0.0687). **E**, Serum levels of HCC-1 in atherosclerosis subjects and atherosclerosis combined with diabetes subjects (AS n=69 vs AS with diabetes n=10, Mann-Whitney test, P=0.8082, SW showed no normal distribution). **F**, Serum levels of HCC-1 in atherosclerosis subjects and atherosclerosis combined with hyperlipidemia subjects (AS n=69 vs AS with hyperlipidemia n=13, Mann-Whitney test, P=0.0177, SW showed no normal distribution).

## DISCUSSION

Atherosclerosis is a chronic inflammatory disease, which is inextricably intertwined with various types of cells, including endothelial cells, or immune cells like monocyte, macrophage, dendritic cell, T cell and so on^5, 27, 28^. As molecular signals inducing leukocytes, chemokines play a pivotal role in inflammation and atherosclerosis by recruiting monocyte into lesions^13^, which is a crucial process in the early stage of atherosclerosis. In the process of plaque progression, pyroptosis plays a major role since a number of pro-inflammatory factors released in this manner of cell death, further recruiting monocytes, exacerbating inflammation and accelerating atherosclerosis^6, 8^. Of note, in addition to recruit monocytes, chemokines are also capable to induce inflammation, cell death like apoptosis or pyroptosis by interacting with cells directly^11, 29, 30^. HCC-1, also named CCL14, is a member of CC-type chemokines and has been studied in hepatocellular carcinoma (HCC) or acute kidney injury (AKI)^16, 17^. However, the function of HCC-1 in inflammation, pyroptosis and atherosclerosis is poorly understood.

To our knowledge, it is the first report of HCC-1 in atherosclerosis. To assess the HCC-1 level in atherosclerotic plaque, We evaluated HCC-1 expression in normal artery tissues and atherosclerotic plaque tissues and our finding revealed that mRNA and protein expression levels of HCC-1 were markedly up-regulated in atherosclerotic plaque tissues, which is confirmed by the result of immunohistochemistry staining analysis. These results suggested that atherosclerotic plaque expressed high levels of HCC-1, which might contribute to plaque formation and progression (Fig. 10).

**Figure 10.** Schematic model of the role of HCC-1 (Hemofiltrate CC chemokine-1) in atherosclerosis. HCC-1 (Hemofiltrate CC chemokine-1) concentration and expression are higher in serum and atherosclerotic plaque in patients with atherosclerosis. HCC-1 secretion enhanced in HUVECs and THP-1 monocytes stimulated by Ox-LDL or LPS in vitro, promoting recruitment of monocytes and M1 polarization, further inducing HUVECs and M1 Macrophages pyroptosis via NF-κB/NLRP3/Caspase-1 pathway.

In atherosclerosis, inflammation and hyperlipidemia are the initiating factors^3, 31^. Because our previous studies showed that atherosclerotic plaque in AS patients had a higher expression of HCC-1, we investigated whether pro-inflammatory stimulus, like LPS and oxidized low-density lipoprotein (Ox-LDL), could induce endothelial cells and Macrophages, two important cell types in atherosclerotic plaque^3, 5^, to enhance the HCC-1 expression. The results indicated that after treated with LPS and Ox-LDL, there was an increase in HCC-1 secretion and gene expression in HUVECs and THP-1 monocytes. Because the immunohistochemistry (IHC) staining showed that endothelial cells expressed high levels of HCC-1, we hypothesized that endothelial cells overexpressing HCC-1 could affect adjaecnt cells. Thus, we made a Co-culture system including HUVECs overexpressing HCC-1, THP-1 monocytes, M0 macrophages or M1 macrophages. The results suggested that HUVECs overexpressing HCC-1 enhanced HCC-1 gene and protein expression in THP-1 monocytes, M0 macrophages and M1 macrophages, indicating that HUVECs overexpressing HCC-1 could further boost HCC-1 expression in THP-1 monocytes and macrophages and then form a positive feedback loop.

In cell experiment, we used recombinant HCC-1 protein to purely imitate the function of increased HCC-1 in patients with atherosclerosis on target cells and lentivirus encoding HCC-1 was used to overexpress HCC-1 in cell lines. As a CC-type chemokine, increased concentration of HCC-1 might induce more monocyte to injured endothelium thus cell adhesion experiment was performed. The results showed that both treatment of recombinant HCC-1 protein and infection of lentivirus encoding HCC-1 in HUVECs or THP-1 monocytes promoted THP-1 monocytes to adhere to HUVECs. To clarify the underlying mechanisms, further studies revealed that both treatment of recombinant HCC-1 protein and infection of lentivirus encoding HCC-1 enhanced ICAM-1、VCAM1 and P-selectin expression in HUVECs and MM7 and MMP10 expression in THP-1 monocytes.

M1 macrophages play a critical role in plaque progression^32^. Na Li et al reported that HCC-1 inhibited colon cancer cell by suppressing M2 polarization of tumor-associated macrophage, while they didn’t evaluate the relationship between HCC-1 and M1 macrophages in tissue and they used an inappropriate concentration of HCC-1 (1μg/ml) to treat M0 macrophages^24^. To clarify the role of HCC-1 in M1 polarization, especially in atherosclerotic plaque, co-localization analysis was performed and our findings demonstrated that in human atherosclerotic plaque, the expression of HCC-1 in Macrophage was co-localized with TNF-α, the specific marker of M1 Macrophage. Further, we used appropriate concentration of HCC-1 (20ng/ml) and HUVECs-Lv-HCC-1 Co-culture system to treat M0 macrophages and the results showed that HCC-1 enhanced TNF-α expression in M0 macrophages, suggesting that HCC-1 promoted the macrophages M1 polarization and might promoted plaque progression.

Endothelial cells and macrophages are two key cell types in atherosclerotic plaque and their inflammation and injury contributes to the formation and progression of atherosclerosis^3, 5^. To investigate whether HCC-1 induced inflammation, our results indicated that recombinant HCC-1 protein, HUVECs-Lv-HCC-1 and co-culture system increased IL-1,IL-6 and TNF-α protein expression both in HUVECs and M1 macrophages. Pyroptosis is a manner of inflammatory cell death and has an important role in cardiovascular disease^6^. Differing from caspase-3 depended apoptosis^33^, classic pyroptosis pathway is caspase-1 depended^7^. After activated, caspase-1 processes the precursor of GSDMD, IL-1β and IL-18 to their mature forms, which leads to the plasma membrane rupture and the following release of inflammatory factors^7^. In this study, we found that HUVECs and M1 Macrophages treated with HCC-1 displayed pyroptotic features, which was showed by increased LDH release and TUNEL and PI positive rate and further confirmed by increased levels of cleaved-IL1β, IL18, ASC, NLRP3, cleaved-caspase-1 and N-GSDMD, indicating that HCC-1 could induce endothelial cells and M1 Macrophages pyroptosis following HCC-1-induced monocytes recruitment and M1 polarization, suggesting that HCC-1 participated in different stages of atherosclerosis.

To investigate the underlying mechanisms, our findings suggested that both recombinant HCC-1 protein and lentivirus encoding HCC-1 boosted TLR4 expression in HUVECs, but not in M1 Macrophages. Because TLR4/NF-κB pathway could initiate pyroptosis^34^ meanwhile NLRP3, the downstream molecule of NF-κB and the upstream molecule of caspase-1, was up-regulated in HUVECs and M1 Macrophages treated with recombinant HCC-1 protein or lentivirus encoding HCC-1, we hypothesized that HCC-1 induced pyroptosis in HUVECs via TLR4/NF-κB/NLRP3/Caspase-1 pathway and in M1 Macrophages via NF-κB/NLRP3/Caspase-1 pathway. The results revealed that TLR4 knocking-down by si-TLR4 transfection blocked the pro-inflammatory and pro-pyroptotic capacity of HCC-1 in HUVECs compared with si-NC transfection group, suggesting that HCC-1 functioned in HUVECs via TLR4/NF-κB/NLRP3/Caspase-1 pathways. Interestingly, although recombinant HCC-1 protein and lentivirus encoding HCC-1 could not enhance TLR4 expression in M1 Macrophages, HUVECs overexpressing HCC-1 boosted TLR4 expression in M1 Macrophages, which might because of the additional secretion of pro-inflammatory factors from HUVECs overexpressing HCC-1.

In animal model, ApoE^−/−^ mice were treated with AAV9-Control and AAV9-HCC-1. Our findings suggested that ApoE^−/−^ mice overexpressing HCC-1 showed a higher average lesion areas of en face, increased atherosclerotic lesions area, lipid deposition, macrophage accumulation and pyroptotic rate and decreased collagen contents in aortic root section. Serum analyses showed that overexpression of HCC-1 increased the concentration of IL-1β and TNF-α in ApoE^−/−^ mice. These results indicated that overexpression of HCC-1 increased inflammatory factors level in serum, induced macrophage accumulation and pyroptosis and then exacerbated atherosclerotic burden. There is a limitation that only male mice were used in this study, with an underlying question that although it is generally believed that males are more prone to atherosclerosis, it is reported that gender is also a biological variable of atherosclerosis, which demonstrates that female mice generally have larger plaques than males in ApoE^−/−^ and LDLR^−/−^ mice^35^. Thus, the function of HCC-1 in female mice requires to be further studied.

It is reported that chemokines could serve as biomarker for diagnosis or prognosis of diseases^36–39^. To assess the potential of HCC-1 serving as a novel biomarker for atherosclerosis, we randomly collected serum from control group and patients with atherosclerosis. Our findings demonstrated that serum level of HCC-1 was remarkably high in patients with atherosclerosis, which was hardly influenced by gender, diabetes or hyperlipidemia. Of note, further studies indicated that serum level of HCC-1 increased with the progression of stenosis, suggesting that HCC-1 might reflect the degree of stenosis. Although HCC-1 showed a high sensitivity in the diagnosis of atherosclerosis, more clinical samples were required to evaluate the specificity of HCC-1 in the diagnosis of atherosclerosis since atherosclerosis tends to occur in the elderly and their health condition is complicated.

Collectively, our findings suggest that chemokine HCC-1 is highly expressed in human atherosclerotic plaque and serum and exacerbates atherosclerosis in vivo. Further studies reveal that HCC-1 promotes the progression of atherosclerosis by recruiting monocyte, inducing M1 polarization and triggering pyroptosis in endothelial cells and M1 Macrophages. Our findings provide strong evidence that HCC-1 is a promising biomarker for the early diagnosis and stratification of atherosclerosis and targeting HCC-1 may serve as a novel preventive or therapeutic approaches in atherosclerosis.

## Acknowledgments

We are thankful to Department of Laboratory Medicine, Nanfang Hospital, Southern Medical University, Guangzhou, People’s Republic of China for providing research facilities. Authors would like to thank Junhui Wang for experiment assistance and Qi Zhang for serum collection.

## Supplemental Material

Major Resources Table Supplemental Figures S1-S9 Table S1

## Highlights

λ HCC-1 concentration in serum is markedly increased in patients with atherosclerosis, which is not influenced by diabetes and hyperglycemia and has the capacity to reflect the degree of stenosis in patients with atherosclerosis.

λ HCC-1 overexpression accelerates the atherosclerotic burden and decreases plaque stability in ApoE^−/−^ mice.

λ HCC-1 overexpression mice have an enhancement in macrophages accumulation, increased pyroptotic rate in ECs and Macrophages in plaque and higher levels of inflammatory factors in serum.

λ LPS and Ox-LDL promote HCC-1 secretion from cultured THP-1 monocytes and ECs.

λ HCC-1 promotes monocytes to adhere to endothelial cells and M1 polarization and induces pyroptosis both in ECs and Macrophages via NF-κB/NLRP3/Caspase-1 pathway.

